# Attentional bias to threat and gray mater volume morphology in high anxious individuals

**DOI:** 10.1101/2020.10.20.346981

**Authors:** Joshua M. Carlson, Lin Fang

**Author notes:** Correspondence should be addressed to: Joshua M. Carlson, Ph.D., Department of Psychological Science, Northern Michigan University, 1401 Presque Isle Avenue, Marquette, MI 49855.

## Abstract

In a sample of highly anxious individuals, the relationship between gray matter volume brain morphology and attentional bias to threat was assessed. Participants performed a dot-probe task of attentional bias to threat and gray matter volume was acquired from whole brain structural T_1_-weighted MRI scans. The results replicate previous findings in unselected samples that elevated attentional bias to threat is linked to greater gray matter volume in the anterior cingulate cortex, middle frontal gyrus, and striatum. In addition, we provide novel evidence that elevated attentional bias to threat is associated with greater gray matter volume in the right posterior parietal cortex, cerebellum, and other distributed regions. Lastly, exploratory analyses provide initial evidence that distinct sub-regions of the right posterior parietal cortex may contribute to attentional bias in a sex-specific manner. Our results illuminate how differences in gray matter volume morphology relate to attentional bias to threat in anxious individuals. This knowledge could inform neurocognitive models of anxiety-related attentional bias to threat and targets of neuroplasticity in anxiety interventions such as attention bias modification.

## Introduction

The human brain is consistently inundated by a multitude of competing sensory inputs, especially within the visual domain. Due to a limited processing capacity, the brain cannot process all sensory input equally and has developed mechanisms by which certain inputs are prioritized for more elaborative processing at the expense of non-prioritized inputs (Desimone & Duncan, 1995). One class of stimuli that are thought to receive prioritized processing are those stimuli that hold biological or emotional significance (Vuilleumier, 2005). This is especially true for emotionally negative or threat-related stimuli that signal the presence of danger (Ohman, Flykt, & Esteves, 2001). The prioritized processing of threat-related stimuli is referred to as a threat bias and the preferential allocation of attentional resources to threatening stimuli is referred to as an attentional bias to threat. This attentional bias to threat appears to occur automatically in the absence of conscious awareness (Carlson & Mujica-Parodi, 2015; Carlson & Reinke, 2008; Fox, 2002; Mogg & Bradley, 2002). Although an attentional bias to threat is adaptive, elevated attentional bias to threat is associated with elevated anxiety (Bar-Haim, Lamy, Pergamin, Bakermans-Kranenburg, & van Ijzendoorn, 2007) and genetic risk for anxiety (Beevers, Gibb, McGeary, & Miller, 2007; Carlson, Mujica-Parodi, Harmon-Jones, & Hajcak, 2012; Fox, Ridgewell, & Ashwin, 2009).

Given the adaptive and maladaptive significance of attentional bias to threat, its underlying neural mechanisms have been extensively studied. Evidence from human lesion and neuroimaging studies indicate that the amygdala plays a critical role in the modulation of spatial attention by threatening stimuli (Anderson & Phelps, 2001; Bach, Hurlemann, & Dolan, 2014; Fu, Taber-Thomas, & Perez-Edgar, 2015; Monk et al., 2008); in part by communicating with visual cortex to prioritize threat-related stimuli (Adolphs, 2004; Amaral & Price, 1984; Carlson, Reinke, & Habib, 2009; Vuilleumier, Richardson, Armony, Driver, & Dolan, 2004). The structural and functional connectivity between the amygdala and areas of the prefrontal cortex (PFC)—such as the anterior cingulate cortex (ACC)—is linked to individual differences in attentional bias to threat such that greater connectivity is associated with elevated attentional bias to threat (Carlson, Cha, Harmon-Jones, Mujica-Parodi, & Hajcak, 2014; Carlson, Cha, & Mujica-Parodi, 2013). The ACC and other PFC regions are also engaged in task-based studies of attentional bias to threat (Armony & Dolan, 2002; Bush, Luu, & Posner, 2000; Fu et al., 2015; Price et al., 2014; White et al., 2016). In addition, greater gray mater volume (GMV) in the basal forebrain/striatum, ACC, and other PFC regions is linked to increased attentional bias to threat (Carlson, Beacher, et al., 2012).

Although research into the neural mechanisms of attentional bias to threat is abundant, the functional significance of individual differences in brain structure has been understudied. To the best of our knowledge, only a single study has explored the relationship between GMV and attentional bias to threat in an unselected sample (Carlson, Beacher, et al., 2012). No studies have explored this relationship in high anxiety individuals—despite the fact that attentional bias to threat is critical in cognitive models of anxiety (Mathews & Mackintosh, 1998) and plays a causal role in the development of anxiety (MacLeod, Rutherford, Campbell, Ebsworthy, & Holker, 2002). Moreover, the efficacy of interventions for anxiety, such as attention bias modification (ABM), are more effectively assessed by tracking changes in brain structure and function following training (Abend et al., 2019; Aday & Carlson, 2017; Britton et al., 2015; Browning, Holmes, Murphy, Goodwin, & Harmer, 2010; Hilland et al., 2020; Hilland, Landro, Harmer, Maglanoc, & Jonassen, 2018; Taylor et al., 2014). Therefore, identifying the neural mechanisms underlying attentional bias to threat will also be important for tracking the neuroplastic effects of ABM and other anxiety interventions.

This study aimed to assess the relationship between brain structure and attention bias behavior in a sample of highly anxious individuals. To this end, participants performed a dot-probe task of attentional bias to threat in a laboratory setting outside the MRI environment. Both traditional attention bias scores and newer trial level bias score measures (Zvielli, Bernstein, & Koster, 2015) from the dot-probe task were utilized. Given the association between elevated anxiety and attentional bias to threat discussed above, participants were selected for heightened levels of attentional bias to threat and anxiety. During a separate testing session, whole brain structural T_1_-weighted MRI scans were acquired. Based on previous research (Carlson, Beacher, et al., 2012), we hypothesized that greater attentional bias to threat would be linked to greater GMV in the ACC and other PFC regions.

## Method

### Participants

One-hundred and eleven right-handed adults (female = 75) between 18 and 38 (*M* = 21.90, *SD* = 4.71) years of age participated in the study. Participants provided written informed consent and received monetary compensation for their participation. The study was approved by the Northern Michigan University Institutional Review Board. Participants included in this report were recruited for a clinical trial assessing the effects of attention bias modification on changes in brain structure (NCT03092609). All data included in this manuscript were collected during the pre-training session. The determination of our sample size was based on the availability of this dataset.

To be included in the study, participants were screened to meet the following criteria (1) right-handed, (2) 18 – 42 years of age, (3) normal (or corrected to normal) vision, (4) no current psychological treatment, (5) no recent history of head injury or loss of consciousness, (6) no current psychoactive medications, (7) not claustrophobic, (8) not pregnant, (9) no metal in the body or other MRI contraindications (10) trait anxiety scores ≥ 40 on the STAI-T (see below for questionnaire information), and (11) attentional bias scores ≥ 7ms in the dot-probe task (see below for task details).

### State-Trait Anxiety Inventory

State and trait anxiety were measured with the Spielberger State-Trait Anxiety Inventory (STAI; Spielberger, Gorsuch, & Lushene, 1970). The STAI-S consists of 20 items and yields a measure of state anxiety (how anxious one currently feels). The STAI-T also consists of 20 items and yields a measure of trait anxiety (how anxious one generally feels). Participants’ STAI-T levels ranged from 40 to 71 (*M* = 51.58, *SD* = 7.31), indicative of a high trait anxious sample.

### Dot-Probe Task

The dot-probe task (MacLeod & Mathews, 1988) was programmed using E-Prime2 (Psychology Software Tools, Pittsburg, PA) and displayed on a 60 Hz 16” LCD computer monitor. Stimuli consisted of 20 fearful and neutral grayscale faces of 10 different actors^1^ (half female; from Gur et al., 2002; Lundqvist, Flykt, & Öhman, 1998) that were cropped to exclude extraneous features. Ratings from a separate sample (*N* = 85) indicate that the fearful faces were perceived as more negative (*M* = 3.83, *SD* = .30) than the neutral faces (*M* = 4.45, *SD* = .52), *t* (18) = 3.23, *p* = .005 (Carlson & Fang, 2020).

Each trial started with a white fixation cue (+) in the center of a black screen for 1000 ms. Two faces were randomly presented simultaneously on the horizontal axis for 100 ms. Participants were seated 59 cm from the screen. Facial stimuli were 5cm × 7cm in size. The target dot appeared at one of the two face locations immediately after the faces disappeared and remained on the screen until a response was made. Responses were recorded with a Chronos E-Prime response box. Participants indicated left-sided targets by pressing the first, leftmost button using their right index finger and indicated right-sided targets by pressing the second button using their right middle finger. Participants were instructed to focus on the central fixation throughout the trial and respond to the target dot as quickly and accurately as possible. The task included five blocks. At the end of each block, participants received feedback about their overall accuracy and reaction times to encourage accurate rapid responses.

The task included congruent trials (dot on the same side as the emotional face), incongruent trials (dot on the same side as the neutral face), and baseline trials (two neutral faces). Faster responses on congruent compared to incongruent trials was considered representative of attentional bias towards the respective emotion. The task consisted of five blocks with 450 trials in total. Each block contained 30 congruent, 30 incongruent, and 30 baseline trials presented in a random order.

### Behavioral Data Preparation and Processing

Data were filtered to include correct responses between 150 and 750 ms post-target onset (Torrence, Wylie, & Carlson, 2017). Attention bias scores were calculated as the mean incongruent – congruent difference in reaction time (in ms). In addition, trial-level bias scores (TLBS) were computed using the steps outlined by Zvielli and colleagues (2015) and Carlson and colleagues (2019). Using these TLBSs, several summary variables were calculated including *Mean_toward_* (Mean of TLBSs > 0 ms; i.e., mean bias towards emotional stimuli), *Mean_away_* (Mean of TLBSs < 0 ms; i.e., mean bias away from emotional stimuli), and variability. TLBS variability was calculated as the summed distance of succeeding TLBS divided by the total number of trial level bias scores. Correlations between attention bias measures can be seen in **Table 1.**

**Table 1.** Correlations between Attention Bias Measures (*N* = 111)

|  | Attention Bias | TLBS Variability | TLBS Toward | TLBS Away |
| --- | --- | --- | --- | --- |
| Attention Bias | -- | .34** | .57** | -.11 |
| TLBS Variability |  | -- | .90** | -.86** |
| TLBS Toward |  |  | -- | -.70** |
| TLBS Away |  |  |  | -- |

Note: TLBS, trial-level bias score. \*\* $p < .001$

### MRI Data Acquisition and Analysis

MRI data were collected with a 1.5 Tesla General Electric whole body scanner^2^ within 1-2 weeks following the behavioral session. High-resolution 3D FSPGR T1-weighted images were obtained using the following acquisition parameters: TR/TI = min/450, flip angle = 9°, FOV = 250, matrix =256 × 256, voxel size = 0.98 × 0.98 mm, slice thickness = 1.2 mm..

We used the automated user-independent voxel-wise measurement approach, known as voxel based morphometry (VBM), to assess the association between regional brain volumes and individual differences in attention bias behavior. The VBM methodology used here is similar to the approach described by Ashburner and Friston (2000). Three-dimensional T_1_-weighted FSPGR MRIs were visually examined for artifacts or abnormalities and manually adjusted to a common origin at the anterior commissure. Images were then pre-processed using established VBM methods in SPM12 (http://www.fil.ion.ucl.ac.uk/spm). Images were first segmented into gray matter (GM), white matter (WM), and cerebrospinal fluid (CSF) tissue types. Segmented gray matter images were normalized to MNI space and smoothed using an 8mm FWHM Gaussian kernel. Measures of intracranial volume (ICV) were obtained from summed global signal of segmented images of GM, WM, and CSF.

Within SPM12 a multiple regression analysis was run which included participants’ (1) Attention Bias, (2) TLBS *Mean_toward_*, (3) TLBS *Mean_away_*, and (4) TLBS Variability scores as predictors of GMV. This regression model also included age and ICV as covariates to control for potential confounding effects on regional gray matter (Ge et al., 2002; Tisserand et al., 2004). We tested for a positive relationship between attention bias measures and GMV. In addition, this model allowed us to compare and contrast different attention bias measures in predicting GMV. An initial whole brain threshold was set to uncorrected *p* < .001 with a 50 voxel cluster threshold. A family-wise error (FWE) small volume correction (SVC) at *p* < .05 was applied to the ACC and other regions of interest (ROIs) previously shown to have GMV correlate with attention bias (i.e., MFG, SFG, IFG, and striatum). The SVC was applied using an 18mm sphere centered on the ROI coordinates reported in Carlson et al (2012). Given recent work exploring sex differences in attentional bias to threat at a behavioral level (Campbell & Muncer, 2017; Carlson, Aday, & Rubin, 2019), exploratory analyses assessing sex differences in the neural correlates of attentional bias are included in the **Supplementary Material**.

## Results

Attentional bias to threat was found to positively correlate with GMV in a number of regions using the initial uncorrected threshold (i.e., *p* < .001, *k* > 50; see **Table 2 & Figure 1**). SVC of ROIs indicated significant associations in the ACC: *t*(104) = 4.02, *k* = 90, xyz = −9, 35, 20, middle frontal gyrus (MFG): *t*(104) = 3.85, *k* = 73, xyz = 29, 59, 5, and striatum: *t*(104) = 3.93, *k* = 127, xyz = 11, 20, −8. Beyond these *a priori* ROIs, whole brain cluster-level FWE associations were observed in the right posterior parietal cortex (rPPC): *t*(104) = 4.11, *k* = 1013, xyz = 56, −53, 38 and bilateral cerebellum: *t*(104) = 4.01, *k* = 943, xyz = −6, −59, −9. No negative associations between attentional bias to threat and GMV were found. In addition, no significant associations were observed for TLBS *Mean_toward_, Mean_away_*, or Variability. However, conjunction analysis between attention bias and TLBS Variability revealed overlapping neural correlates (see **Supplementary Table 2**) and a direct comparison of attention bias scores with TLBSs (i.e., attention bias > TLBS) did not result in any significant differences. Finally, results of exlporatory analyses of sex differences in the neural correlates of attentional bias to threat can be found in the **Supplementary Materials**.

**Figure 1.**
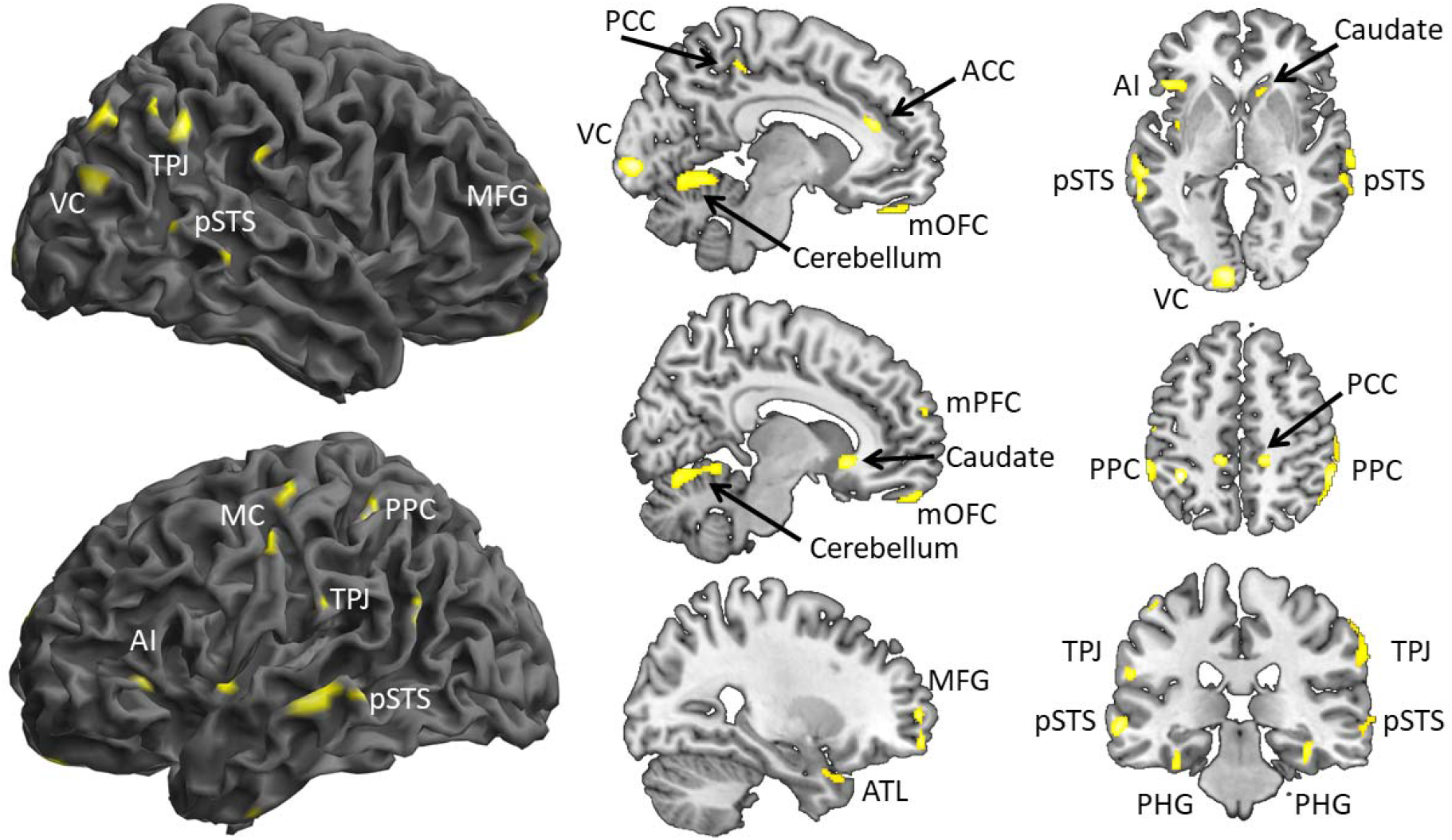
Attentional bias to threat correlated with regional gray matter volume (GMV) in the anterior cingulate cortex (ACC), anterior insula (AI), anterior temporal lobe (ATL), caudate, cerebellum, medial orbitofrontal cortex (mOFC), middle frontal gyrus (MFG), motor cortex (MC), parahippocampal gyrus (PHG), posterior cingulate cortex (PCC), posterior parietal cortex (PPC), posterior superior temporal sulcus (pSTS), temperoparietal junction (TPJ), and regions of the visual cortex (VC). Greater GMV was associated with greater attentional bias to threat. Images thresholded at uncorrected *p* <.001, k> 50 voxels.

**Table 2:** GMV Correlates of Attentional Bias to Threat

| Region | Hemisphere | MNI Coordinates |  |  | Voxels | Peak<br>t value | Sig. |
| --- | --- | --- | --- | --- | --- | --- | --- |
|  |  | X | Y | Z |  |  |  |
| *Parietal Cortex* | L | -56 | -41 | 53 | 155 | 4.05 | * |
|  | L | -38 | -47 | 50 | 144 | 4.54 | * |
|  | L | -60 | -27 | 24 | 55 | 3.76 | * |
|  | R | 56 | -53 | 38 | 1013 | 4.11 | *** |
| *Cerebellum* | L-R | -6 | -59 | -9 | 943 | 4.01 | *** |
|  | L | -21 | -45 | -57 | 545 | 4.26 | * |
|  | L | -35 | -75 | -47 | 137 | 3.56 | * |
| *Anterior Cingulate Cortex* | L | -9 | 35 | 20 | 90 | 4.02 | ** |
| *Posterior Cingulate Cortex* | L | -12 | -36 | 50 | 104 | 3.96 | * |
|  | R | 15 | -38 | 54 | 130 | 4.33 | * |
|  | L-R | 3 | -60 | 27 | 88 | 3.50 | * |
| *Temporal Parietal Junction* | L | -53 | -62 | 21 | 169 | 4.53 | * |
|  | R | 62 | -57 | 14 | 92 | 3.57 | * |
| *Superior Temporal Sulcus* | L | -63 | -42 | -2 | 379 | 4.22 | * |
|  | R | 71 | -23 | -2 | 258 | 3.65 | * |
| *Visual Cortex* | L | -9 | -95 | -3 | 388 | 4.28 | * |
|  | R | 42 | -80 | 21 | 130 | 4.18 | * |
|  | R | 32 | -74 | 44 | 197 | 3.76 | * |
| *Striatum* | R | 11 | 20 | -8 | 192 | 3.93 | ** |
| *Middle Frontal Gyrus* | R | 29 | 60 | -9 | 112 | 3.92 | * |
|  | R | 29 | 59 | 5 | 73 | 3.85 | ** |
|  | R | 32 | 56 | 23 | 57 | 3.48 | * |
| *Anterior Insula* | L | -47 | 26 | -2 | 149 | 3.88 | * |
| *Posterior Insula* | L | -45 | -3 | 9 | 175 | 3.53 | * |
| *Parahippocampal Gyrus* | L | -36 | -23 | -27 | 203 | 3.85 | * |
|  | R | 36 | -27 | -18 | 97 | 3.72 | * |
| *Medial Prefrontal Cortex* | R | 14 | 65 | 20 | 71 | 3.82 | * |
| *Superior Frontal Gyrus* | L | -21 | 12 | 68 | 56 | 3.78 | * |
| *Medial Orbitofrontal Cortex* | L | -8 | 47 | -26 | 105 | 3.65 | * |
|  | R | 8 | 57 | -24 | 97 | 3.72 | * |
| *Anterior Temporal Lobe* | L | -53 | -3 | -39 | 72 | 3.51 | * |
|  | R | 24 | 15 | -32 | 82 | 3.55 | * |
| *Precentral Gyrus* | L | -54 | -18 | 53 | 50 | 3.53 | * |
|  | L | -41 | -20 | 60 | 71 | 3.5 | * |

\* $p < .001$ uncorrected voxel-level, \*\* $p < .05$ SVC cluster-level, \*\*\* $p < .05$ FWE cluster-level

## Discussion

In a sample of high trait anxious individuals, we found that individual differences in attentional bias to threat correlated with regional differences in GMV. Consistent with previous research in unselected samples (Carlson, Beacher, et al., 2012), cluster-level FWE corrected ROI analyses suggest that greater GMV in the ACC, MFG, and basal forebrain/striatum are linked to elevated attentional bias to threat. Furthermore, in our high anxiety sample, whole brain FWE corrected analyses indicate that elevated attentional bias to threat is additionally associated with greater GMV in the rPPC and cerebellum. A number of additional regions displayed this same association at an uncorrected level (see **Table 2**). No regions showed the opposite (i.e., negative) association. In our exploratory analyses, we observed sex differences in distinction sub-regions of the rPPC. In a relatively more medial and dorsal cluster, there was a negative correlation between GMV and attentional bias in males, but not females. On the other hand, in a relatively more lateral and ventral cluster, there was a negative association between GMV and attentional bias in females, but not males (see **Supplementary Material** for more information). In sum, we present novel evidence that GMV in distributed brain regions such as the ACC, MFG, striatum, rPPC, and cerebellum (among other regions) is linked to attentional bias.

The associations between GMV and attentional bias to threat observed here builds on earlier research in unselected samples (Carlson, Beacher, et al., 2012). Consistent with this previous work, we found that greater GMV in the ACC, MFG, and basal forebrain/striatum was related to greater attentional bias to threat. Not only did this earlier research use an unselected sample, but also used a backward masking procedure to restrict the processing of the fearful face cues. Thus, our findings generalize the association between attentional bias to threat and GMV in the ACC (among other regions) to high anxiety individuals and unmasked threat signals afforded more elaborative processing. The ACC is thought to play a role in conflict monitoring and resolution (Botvinick, Cohen, & Carter, 2004; Botvinick, Nystrom, Fissell, Carter, & Cohen, 1999). In the context of attentional bias to threat, the ACC may detect conflict between the location of the threat-related stimulus and the goal-relevant target stimulus (i.e., incongruent trial types; Fu et al., 2015; Price et al., 2014). Indeed, conflict related ERPs (i.e., N2) are elicited in the dot-probe task (Andrzejewski & Carlson, 2020). When conflict is detected, the ACC and other PFC regions (e.g., MFG) may regulate the duration of attentional focus on the threat stimulus as well as the eventual disengagement and reengagement of attention to the goal-relevant target stimulus.

In addition to these previously implicated regions, novel associations between attentional bias to threat and GMV were observed in the rPPC and bilateral cerebellum. The rPPC is traditionally thought to play a role in spatial cognition including spatial attention (Culham & Kanwisher, 2001). It is part of a frontoparietal network involved in attentional control (Scolari, Seidl-Rathkopf, & Kastner, 2015). Interestingly, patients with lesions to the rPPC generally neglect (or fail to attend to) visual stimuli in the contralateral hemi-field, but do not neglect emotional or threat-related stimuli in the contra-lateral hemi-field (Vuilleumier & Schwartz, 2001). Suggesting that the rPPC is involved in spatial attention, but is not necessary for emotional attention. Yet, although not necessary, structural variability in this structure appears to be linked to individual differences in attentional bias to threat in high anxiety individuals. In addition, the cerebellum has recently been implicated in anxiety-related symptoms and behaviors (Moreno-Rius, 2018). Cerebellar GMV differences and hyperactivity to threat-related stimuli have been observed in a variety of anxiety disorders (Moreno-Rius, 2018). Our results suggest that it is also involved in attentional bias to threat. Consistent with this notion, many cerebellar sub-regions are functional connected to the amygdala, ACC, PPC, and other regions involved in the allocation of spatial attention to threat (O’Reilly, Beckmann, Tomassini, Ramnani, & Johansen-Berg, 2010; Sang et al., 2012).

At an uncorrected threshold, we observed more widespread associations with attentional bias to threat with GMV in regions of the PFC, temporal lobes, and visual cortex. Many of these structures, including the ACC, insula, temperoparietal junction (TPJ), posterior superior temporal sulcus (pSTS), parietal cortex, and visual processing regions reveal patterns of intrinsic functional connectivity with the amygdala that are linked to attentional bias to threat and structural connectivity (Carlson et al., 2013). That is, greater amygdala connectivity in these regions has been linked to greater attentional biases to threat. However, consistent with previous research (Carlson, Beacher, et al., 2012), we did not find an association between amygdala GMV and attention bias. Evidence from animal models indicates that differences in VBM measures of GMV are linked to dendritic spine density (Keifer et al., 2015). Given that increased dendritic spine density is associated with fear learning and represents the capacity for afferent signals into a structure, differences in GMV may reflect the connectivity or processing capacity of a region. If this is indeed true, it may explain why we do not see GMV differences in the amygdala. That is, differences in ACC, insula, TPJ, pSTS, parietal cortex, and visual cortical GMV may (at least in part) be due to the number of synapses from amygdala efferents. However, this hypothesis remains untested and future research is needed to assess its validity.

We included both traditional and newer measures of attentional bias. However, the newer TLBS measures of attentional bias did not correlate with GMV (at *p* < .001, *k* > 50). These null effects are likely reflective of similar, but weaker effects in these measures. Indeed, conjunction analysis indicates some overlap in GMV correlates between TLBS variability and the traditional measure of attentional bias (see **Supplementary Table 2**). In addition, the traditional attention bias > TLBS measures contrast in SPM did not reveal any significant differences. Indeed, attentional bias was moderately correlated with TLBS variability and TLBS Mean Positive, but all TLBS measures were highly correlated (see **Table 1**). Thus, TLBS measures are related to attentional bias, but appear to be more related to variability and this additional component may result in weaker associations with GMV.

Our results implicate a widespread network of regions where brain morphology is related to an anxiety-related symptom—attentional bias to threat. It should be emphasized that the direction of this relationship is unclear. That, is variability in regional GMV in this network may be an underlying risk factor for elevated attentional bias to threat, which has been shown to be a causal mechanism for the development of anxiety (MacLeod et al., 2002). Or, these GMV differences may be the consequence of this behavior. Future longitudinal research will be needed to test these two possibilities. It has been hypothesized that tracking changes in GMV following anxiety reducing interventions such as ABM may be an effective way to gauge the success of the treatment in reorganizing the brain (Aday & Carlson, 2017). Indeed, recent research suggests that ABM produces structural changes in the brain (Abend et al., 2019). The regions identified in the current report (see **Table 2**) may be utilized as a priori ROIs for future studies assessing neuroplasticity in ABM. In addition, GMV in these regions could potentially be used to predict who is most likely to benefit from such treatments. That is, understanding how brain structure relates to attentional bias across individuals could be essential for understanding how ABM may induce neuroplasticity.

### Limitations, Strengths, and Conclusions

Like any study, this study had several strengths and weaknesses. A strength of our study is the relatively large sample size. However, this sample included more female than male participants and was comprised of primarily younger adults. Future research is needed to determine if the effects observed here generalize to younger and older age groups. In addition, given the well-established association between elevated anxiety and attentional bias to threat (Bar-Haim et al., 2007), we selected participants for heightened levels of attentional bias to threat and anxiety. On one hand, the composition of our sample is a strength as the relationship between GMV and attentional bias is highly relevant in neurocognitive models of anxiety. On the other hand, by selecting only individuals at the higher ends of each spectrum, we limited the range/variability of our measures, which precludes us from generalizing these findings to low anxiety individuals with low levels of attentional bias. Although it should be noted that many of our findings are consistent with earlier work using an unselected sample (Carlson, Beacher, et al., 2012). Nevertheless, future research should explore the association tested here across a broader spectrum of anxiety levels.

In conclusion, we found that greater GMV in the ACC, MFG, and basal forebrain/striatum is linked to greater attentional bias to threat in high anxiety individuals, which replicates such associations in unselected samples (Carlson, Beacher, et al., 2012). We provide additional novel evidence that elevated attentional bias to threat in high trait anxious individuals is associated with greater GMV in the rPPC, cerebellum, and a number of other regions (see **Table 2**). Exploratory analyses provide initial evidence that distinct sub-regions of the rPPC may contribute to attentional bias to threat differently across the sexes (see **Supplementary Material**). Collectively, these findings add to our understanding of how differences in brain structure relate to attentional bias to threat in anxious individuals. Such knowledge may be informative for neurocognitive models of attentional bias in anxiety and for tracking neuroplasticity in interventions (e.g., ABM) for anxiety.

## Acknowledgements

Research reported in this publication was supported by the National Institute of Mental Health of the National Institutes of Health under Award Number R15MH110951. The content is solely the responsibility of the authors and does not necessarily represent the official views of the National Institutes of Health. We would like to thank Jeremy Andrzejewski, Taylor Susa, Katie Elwell, and all the other students in the Cognitive × Affective Behavior & Integrative Neuroscience (CABIN) Lab at Northern Michigan University as well as Rochelle Milano and Annette Gustafson at UPHS-Marquette for assisting in the collection of this data.

## Disclosures

### Ethics approval

The study was approved by the Northern Michigan University Institutional Review Board (HS13- 555).

### Consent to participate

All participants provided written informed consent.

### Consent for publication

NA

### Availability of data and material

Data available upon request. Data will be available on the NIMH Data Archive 9/15/2021.

### Code availability

NA

### Open Practice Statement

The data will be available on the NIMH Data Archive 9/15/2021. The larger study from which this data comes from is registered at clinicaltrials.gov under NCT03092609.

## Footnotes

1 Fearful and Neutral face stimuli were from actors: 207, 208, 213, and 217 (Gur et al., 2002) as well as AF14, AF19, AF22, AM10, AM22, AM34 (Lundqvist, Flykt, & Öhman, 1998).

2 MRI data were collected on two identical scanners (scanner 1 *n* = 64 & scanner 2 *n* = 47). A comparison of whole brain GMV (*p* = .86) and ICV (*p* = .55) across scanners indicated no systematic differences in scanning location.

