## Supplementary Material for "Attentional bias to threat and gray mater volume morphology in high anxious individuals"

**Analyses exploring the moderation of the association between GMV and attention bias by sex**

Variability in the gray mater volume (GMV) of the basal forebrain/striatum, anterior cingulate cortex (ACC), and other prefrontal cortex (PFC) regions has been linked to individual differences in attentional bias to threat (Carlson et al., 2012). At a behavioral level, recent research has explored the possibility of sex differences in attentional bias to threat (Campbell & Muncer, 2017; Carlson, Aday, & Rubin, 2019). In addition, sex differences in GMV have been linked to a number of affective traits (Carlson, Depetro, Maxwell, Harmon-Jones, & Hajcak, 2015; Welborn et al., 2009) and there are well established sex differences in the prevalence rates of anxiety (Kessler, Petukhova, Sampson, Zaslavsky, & Wittchen, 2012), which may be related to differences in brain morphology. However, no studies have assessed the role of sex in moderating the relationship between GMV and attentional bias to threat. Therefore, in an exploratory analysis we tested for a moderating role of participant sex, but did not have specific hypotheses for this exploratory analysis.

The supplementary regression analysis included participants' (1) Attention Bias Measure^[1]^, (2) Sex, and (3) Attention Bias x Sex Interaction Terms as predictors of GMV as well as age and ICV as covariates. In this analysis, we assessed the moderating role of sex by evaluating associations that were (1) greater in females compared to males or (2) greater in males compared to females. A whole brain threshold was set to uncorrected *p* < .001 with a 50 voxel cluster threshold. Note that male and female participants did not differ in age, anxiety, or attention bias measures (see **Supplementary Table 1**).

**Supplementary Table 1:** Means (Standard Deviations) of Behavioral Measures by Sex

| **Factor** | **Female**  **n = 75** | **Male**  **n = 36** | **Stats** |
| --- | --- | --- | --- |
| **Age** | 21.51 (4.72) | 22.72 (4.63) | *t*(109) = 1.28, *p =* .20 |
| **STAI-T** | 51.69 (7.80) | 51.33 (6.25) | *t*(109) = -0.24, *p =* .81 |
| **Attention Bias** | 14.91 (9.07) | 16.36 (8.94) | *t*(109) = 0.79, *p =* .43 |
| **TLBS Variability** | 44.79 (11.57) | 42.26 (10.03) | *t*(109) = -1.12 , *p =* .26 |
| **TLBS Mean Toward** | 60.41 (17.25) | 56.16 (13.99) | *t*(109) = -1.30, *p =* .20 |
| **TLBS Mean Away** | -50.97 (14.87) | -48.22 (14.04) | *t*(109) = 0.93, *p =* .36 |

*Note*: STAI, Spielberger State-Trait Anxiety Inventory; TLBS, trial-level bias score.

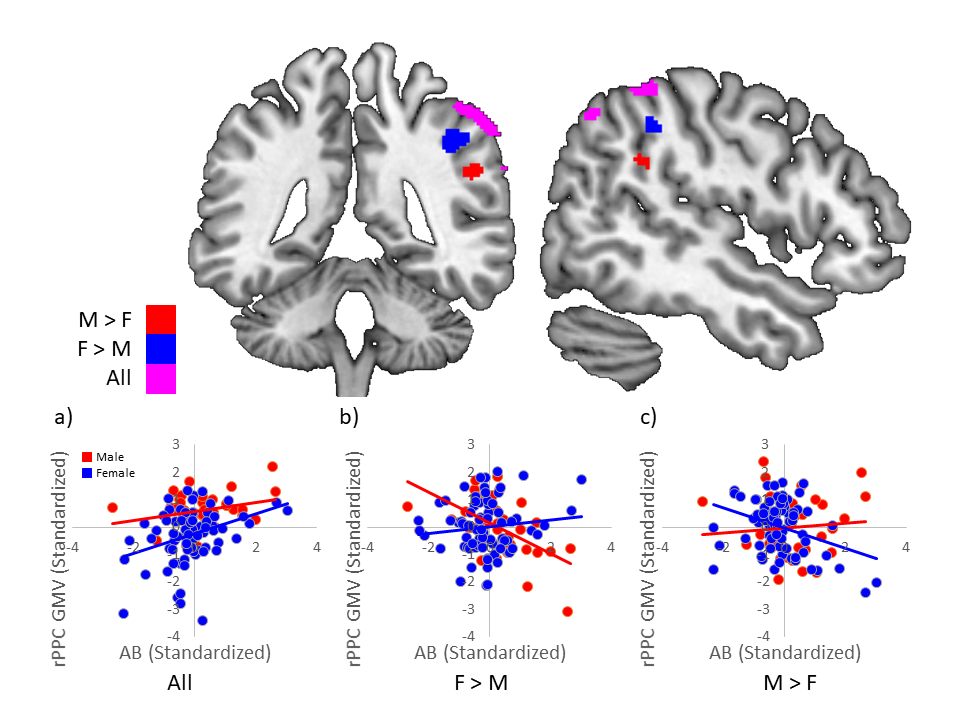

***Supplementary Figure.*** *Three distinct regions of the right posterior parietal cortex where gray matter volume (GMV) differentially correlated with attentional bias to threat. (a) In a more superior superficial region, GMV was positively correlated with attentional bias in both males (M) and females (F). (b) In the blue region, there was a significant negative correlation in males (r = -.55, p < .001), but no association in females (r = .11, p = .17). (c) In the red region, there was a significant negative correlation in females (r = -.35, p < .001), but no association in males (r = .09, p = .30). Note “All” refers to both males and females.*

**Supplementary Results and Discussion of Sex Differences**

SVC centered on the rPPC cluster implicated in Model 1 revealed distinct clusters within the rPPC where the association between GMV and attentional bias was (1) greater in females (compared to males): *t*(104) = 4.53, *k* = 232, xyz = 44, -41, 42 and (2) greater in males compared to females: *t*(104) = 3.57, *k* = 70, xyz = 51, -44, 30 (see **Supplementary Figure**). Closer examination revealed that these differential associations were driven by negative correlations in males (*r* = -.55, *p* < .001, but not females *r* = .11, *p* = .17) at xyz = 44, -41, 42 and females (*r* = -.35, *p* < .001, but not males *r* = .09*, p* = .30) at xyz = 51, -44, 30. That is, for the contrast female attention bias > male attention bias, “greater” indicates the relationship we less negative. The same was true for the male attention bias > female attention bias contrast. These two clusters were the only associations to survive the threshold of *p* < .001, *k* > 50.

We present initial evidence for sex-specific associations with attentional bias in discrete sub-regions of the rPPC. In males, there was a negative relationship between GMV and attentional bias in a relatively more medial and dorsal cluster (see **Supplementary Figure**). In females, there was a negative relationship between GMV and attentional bias in a relatively more lateral and ventral cluster (see **Supplementary Figure**). Both of these regions are discrete from a third cluster that had a positive relationship with attentional bias to threat in our primary analysis. Although we did not have specific hypotheses about this effect, sex differences in GMV have been observed in the rPPC and these differences are linked to spatial cognition (i.e., mental rotation; Koscik, O'Leary, Moser, Andreasen, & Nopoulos, 2009). Thus, sex differences in rPPC morphology appear to be related to aspects of spatial cognition, including spatial attention. It should be noted, however, that female and male participants in the current study did not differ in their attentional bias to threat (see **Supplementary Table 1**). Yet, it appears that the underlying structural correlates of attentional bias in the rPPC differ between males and females. Given that GMV appears to be reflective of dendritic spine density (Keifer et al., 2015), this finding may indicate differing locations for inputs relevant (or antagonistic) to the spatial processing/coding of environmental threats between males and females. Although we provide initial evidence for sex differences in the rPPC related to attentional bias to threat, future research will be needed to corroborate these sex-specific associations.

**Supplementary Table 2:** Conjunction Analysis of Attention Bias and TLBS Variability on GMV

|  |  | **MNI Coordinates** | | |  | **Peak** | **Peak** |
| --- | --- | --- | --- | --- | --- | --- | --- |
| **Region** | **Hemisphere** | **X** | **Y** | **Z** | **Voxels** | ***t* value** | ***p* value** |
| *Cerebellum* | L | -21 | -44 | -53 | 368 | 3.00 | .002 |
|  | R | 3 | -63 | -47 | 49 | 2.74 | .004 |
|  | R | 11 | -65 | -8 | 27 | 2.63 | .005 |
| *Visual Cortex* | L | -23 | -99 | -2 | 39 | 2.63 | .005 |
|  | L | -11 | -93 | -5 | 45 | 2.55 | .006 |
|  | R | 14 | -90 | 27 | 102 | 3.07 | .001 |
| *Middle Frontal Gyrus* | L | -38 | 53 | 8 | 38 | 2.94 | .002 |
| *Parahippocamal Gyrus* | R | 35 | -30 | -24 | 46 | 2.86 | .003 |
| *Anterior Temporal Lobe* | R | 36 | 21 | -27 | 56 | 2.79 | .003 |
| *Inferior temporal Gyrus* | L | -59 | -38 | -26 | 37 | 2.61 | .005 |
|  | R | 56 | -12 | -38 | 44 | 2.77 | .003 |

*Note*. This conjunction analysis used a threshold of *p* < .01 uncorrected and *k* > 25 voxels. TLBS, trial-level bias score.

1. Note that we used the traditional attention bias measure in this exploratory analysis as (1) it was the measure to most strongly correlate with GMV measures in our primary analysis and (2) contrasts between the traditional measure and trial-level bias score measures did not find significant differences between the measures.
